# EnzymeTuning: a Deep-learning-based Toolbox for Optimizing Enzyme-constrained Metabolic Modeling with Enhanced Proteome Abundance Prediction

**DOI:** 10.1101/2025.08.23.670849

**Authors:** Xueting Wang, Yingping Zhuang, Guan Wang, Hongzhong Lu

**Affiliations:** State Key Laboratory of Bioreactor Engineering, East China University of Science and Technology, Shanghai 200237, P. R. China; State Key Laboratory of Microbial Metabolism, School of Life Science and Biotechnology, Shanghai Jiao Tong University, Shanghai 200240, P. R. China

**Keywords:** generative adversarial networks, proteome abundance prediction, *kcat* optimization

## Abstract

The accuracy of enzyme kinetic parameters, particularly the turnover number (*k*cat), is critical for the predictive power of enzyme-constrained genome-scale metabolic models (ecGEMs). However, current kinetic datasets remain sparse and often fail to capture *in vivo* enzyme behavior, compromising model predictive capacity. To address these challenges, we developed EnzymeTuning——a generative adversarial network (GAN)-based framework for the global *kcat* optimization. This approach significantly enhances both the accuracy and proteome-level coverage of ecGEM predictions. Moreover, by incorporating literature-derived protein degradation constants (*kdeg*), we inferred the protein synthesis rate and systematically evaluated their impact on model performance. The framework was validated across distinct yeast species, including *Saccharomyces cerevisiae*, *Kluyveromyces lactis*, *Kluyveromyces marxianus*, and *Yarrowia lipolytica*, demonstrating its generalizability. Further, we found that EnzymeTuning facilitates the identification of context-specific enzyme usage patterns and adaptive catalytic resource allocation under varying carbon-to-nitrogen (C/N) ratios, showcasing the substantial potential of our toolbox for integrative omics analysis. Overall, EnzymeTuning provides a robust and scalable solution for refining kinetic parameters in ecGEMs, thereby promoting the wide applications of these computational models in systems and synthetic biological studies.

## Introduction

Enzyme-constrained genome-scale metabolic models (ecGEMs) have emerged as powerful computational platforms for simulating cellular growth, metabolic reprogramming, and protein resource allocation. By integrating enzymatic capacity constraints, ecGEMs provide unprecedented insights into the regulation of metabolic networks and facilitate the rational optimization of microbial biosynthesis processes, particularly for the production of high-value compounds such as biofuels and pharmaceutical precursors [1, 2]. Compared with traditional GEMs, ecGEMs more accurately account for the enzymatic limitations, thereby enhancing predictive precision. They have been widely applied in metabolic engineering to enhance the yield of target products [3]. Notably, the development of the GECKO 3.0 toolbox has substantially streamlined ecGEM construction and implementation across diverse species, enabling applications in both microbial systems (e.g., *Saccharomyces cerevisiae*) and mammalian systems (e.g., human cells) for metabolic optimization [4]. Central to the predictive capacity of ecGEMs is the accurate parameterization of enzyme kinetics, particularly the enzyme turnover number (*kcat*) [1, 5]. In metabolic networks, *kcat* values not only directly determine reaction fluxes but also modulate global metabolic flux distribution through protein resource allocation [5]. Nevertheless, the experimental determination of physiologically relevant *kcat* values remains a persistent methodological challenge. Conventional *in vitro* enzyme activity assays require protein purification under non-physiological conditions, which are both time- and resource-intensive. Moreover, these measurements often lack biological relevance due to environmental discrepancies (e.g., pH, temperature) [6, 7]. The inherent low-throughput of such assays has further resulted in sparse and error-prone kinetic datasets, thereby limiting the application of ecGEMs [8].

To overcome these limitations, deep learning (DL)-based approaches for *kcat* parameter tuning have emerged as promising tools for large-scale kinetic parameter optimization [9, 10]. For instance, DLKcat integrated graph neural networks (GNNs), convolutional neural networks (CNNs), and Bayesian inference to predict *kcat* values across 332 yeast species, improving both the proteome-level coverage of ecGEMs and the accuracy of phenotype predictions [11]. Nonetheless, DLKcat exhibited limited performance in protein abundance prediction, as indicated by modest Pearson correlation coefficients (PCC) (∼0.3) between predicted and measured abundances [11]. Recently, Kroll et al. developed TurNup, which leveraged Transformer architectures with self-attention mechanisms to decode long-range dependencies within enzyme sequences, thereby effectively improving protein allocation predictions [12]. TurNuP outperformed previous CNN-based models in *kcat* prediction accuracy. However, both DLkcat and TurNuP remain heavily dependent on protein sequence information with their predictive accuracy decreasing notably for enzymes exhibiting low sequence similarity to the training set or lacking well-annotated protein sequences.

Generative adversarial networks (GANs)—a deep learning algorithm based on adversarial training between a generator and a discriminator—have been widely applied in diverse domains, including biological engineering and biomedicine [13, 14]. For instance, GANs have been employed to generate novel molecular structures, accelerating screening and optimization in drug discovery and molecular design [15]. In biomedical image processing, GANs have improved image quality through tasks such as super-resolution, reconstruction, and segmentation, leading to enhanced diagnostic accuracy [16]. Additionally, GAN-based models have facilitated dimensionality reduction and feature extraction in single-cell RNA sequencing analysis, facilitating deeper insights into cellular heterogeneity [17]. Unlike sequence-dependent approaches such as CNNs and Transformers, GANs offer a data-efficient and distribution-oriented strategy for kinetic parameter inference. Recent studies have introduced GAN variants for kinetic parameter tuning in metabolic models, allowing the exploration of physiological metabolic states with limited data and reduced computational requirements [18]. Nevertheless, the application of GANs to enzyme kinetic parameter inference with ecGEMs remains underexplored. Moreover, conventional GAN methods often require large datasets and substantial computational resources, posing challenges for genome-scale metabolic models. Therefore, the development of a GAN-based framework capable of efficiently addressing high-dimensional enzyme kinetic parameter spaces for the global optimization of ecGEMs represents an important and timely research direction.

In this study, we developed a GAN-based framework named EnzymeTuning that enables global optimization of *kcat* parameters in ecGEMs. By leveraging adversarial training, EnzymeTuning can generate realistic *kcat* distributions, thus overcoming the limitations associated with enzymes lacking homologous sequences. This approach effectively improved the quantitative prediction of protein abundances in ecGEMs. Beyond *kcat* inference, EnzymeTuning was extended to estimate protein synthesis rates and to assess the predictive capacity of ecGEMs when such rates were incorporated as constraints. We validated the general applicability and superiority of EnzymeTuning by optimizing ecGEMs for *Kluyveromyces lactis*, *Kluyveromyces marxianus*, and *Yarrowia lipolytica*, achieving substantial improvements in protein abundance prediction. Furthermore, to investigate context-specific enzyme utilization, we analyzed *in vivo kcat* distributions under varying carbon-to-nitrogen (C/N) ratios, thereby elucidating adaptive protein resource allocation strategies. Collectively, our study introduces a robust, scalable, and generalizable computational framework for refining kinetic parameters in ecGEMs, paving the way for improved predictive modeling in systems and synthetic biology.

## Results

### Framework of EnzymeTuning

The EnzymeTuning framework employed a three-step optimization workflow initialized with Bayesian posterior distributions of *kcat* values, as illustrated in **Figure 1** [11]. In Step 1, high-performance parameter sets were identified based on their ability to accurately predict both growth phenotypes and quantitative protein abundances, forming a labeled training dataset for a conditional generative adversarial network (cGAN) [20]. In Step 2, the cGAN architecture was iteratively trained to learn and reproduce the distribution of high-performance *kcat* sets, using performance-based labels (high-performance/low-performance) as guidance. After each training iteration, the top 100 parameter sets were validated with the ecGEM. In Step 3, the generated parameter sets were systematically evaluated under various cultivation conditions, including carbon-limited and nitrogen-limited conditions, conditions with alternative nitrogen source (e.g. glutamate, iso-leucine, and phenylalanine), and batch cultivations of *S. cerevisiae*, to assess both their rationality and the generalizability of the framework. In Step 1, the ecGEM of *S. cerevisiae* was updated using the *kcat* parameter set derived from the Bayesian posterior distribution (**Figure S1**). The root mean squared error (RMSE) between measured and predicted values under different conditions including batch cultivations and carbon-limited continuous conditions was used as the evaluation metric for growth phenotype prediction. After 100 generations, the ecGEM parameterized with the posterior *kcat* values achieved RMSEs below 1, and the *kcat* distribution closely resembled that obtained via deep-learning-based methods (**Figures S1a, S1b**). Moreover, the updated ecGEM successfully reproduced the Crabtree effect in *S. cerevisiae*, thereby improving the performance of phenotype prediction compared with the previous version (**Figure S1c**). For quantitative protein abundance prediction, the predicted abundances of metabolic enzymes in the updated ecGEM were compared with experimentally measured proteomics data under carbon-limited conditions (**Figure S1d**). The average Pearson correlation coefficient (PCC) between predicted and measured values was 0.318 (*P* value <0.05), indicating a statistically significant but modest predictive performance. Relative to the proteomics dataset, the updated ecGEM exhibited a high proportion of extreme enzyme abundance predictions—defined as predicted/measured abundance ratios greater than 100 or less than 0.01—which accounted for approximately one-fourth to one-third of all predictable enzymes. The combination of a modest PCC and a substantial fraction of extreme predictions further indicated the suboptimal accuracy of protein abundance prediction (**Figures S1e, S1f**). Subsequently, we evaluated the performance of the GAN framework for global *kcat* parameter tuning. Given that the ecGEM parameterized with Bayesian-posterior *kcat* values already achieved accurate growth phenotype predictions, our optimization primarily focused on *kcat* parameters that were insensitive to growth phenotype prediction with poor accuracy in protein abundance prediction. In this study, we conducted a global sensitivity analysis of *kcat* parameters using the SALib method (**Figure 2a**) [19]. Specifically, each *kcat* value was individually perturbed by random sampling within five ranges of the initial *kcat* values: 10%–30%, 50%–70%, 70%–90%, 90%–110%, and 150%–170%. The first-order sensitivity coefficient (S1) and the total-order sensitivity coefficient (ST) from SALib were used to quantify parameter sensitivity (**Figures 2b, 2c**). Since higher S1 and ST values indicated greater parameter influence on growth phenotype prediction, the top 25% of *kca*t parameters with the highest S1 and ST values were classified as sensitive. Here, S1 quantified the effect of individual *kcat* variations, whereas the ST value accounted for the combined influence of multiple *kcat* parameters. By intersecting the set of *kcat* parameters that were insensitive to growth phenotype prediction with those associated with extreme protein abundance predictions (**Figures 2d–f**), a total of 237 *kcat* parameters were identified for subsequent optimization using the EnzymeTuning framework.

**Figure 1.**
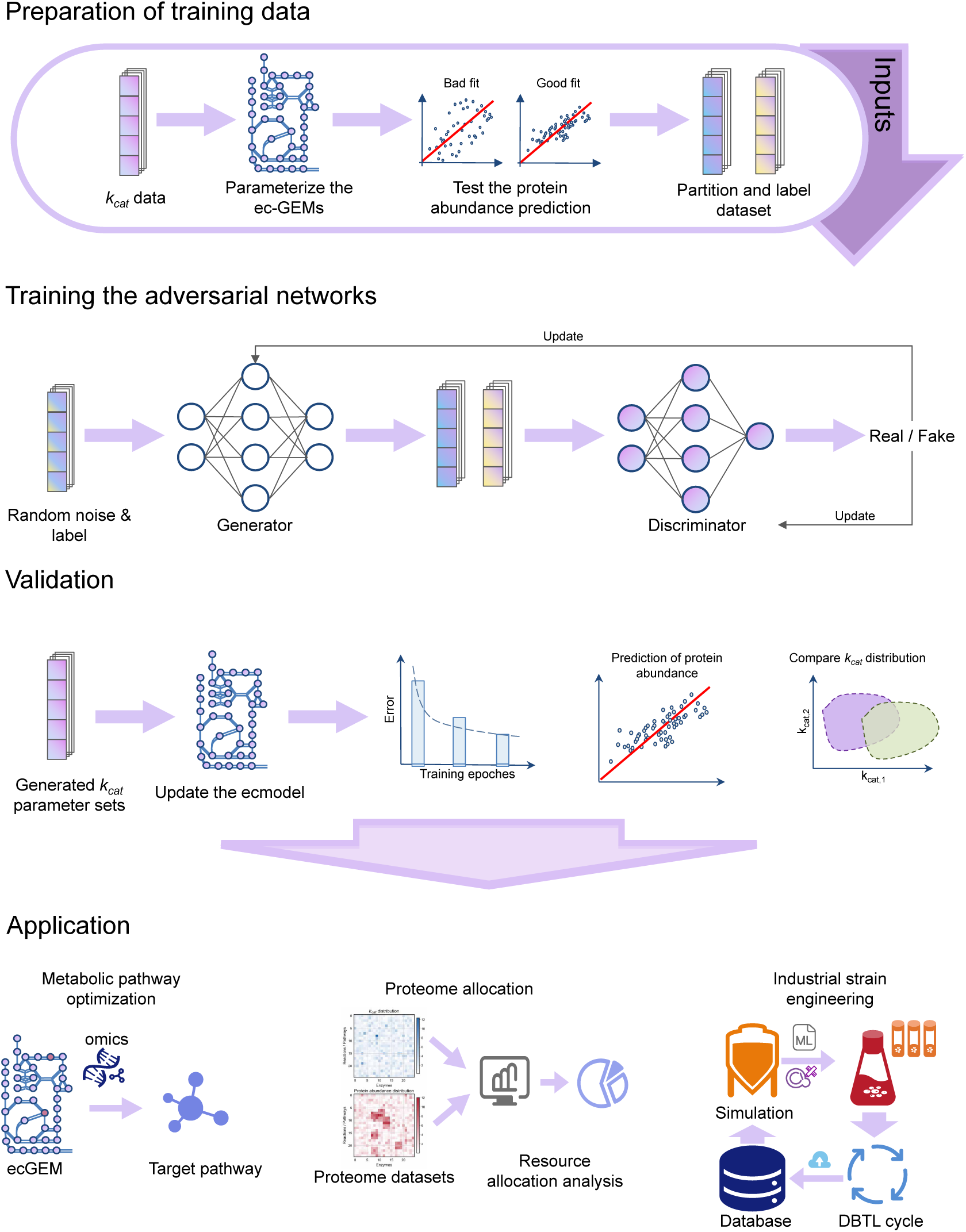
Overview of the EnzymeTuning framework.

**Figure 2.**
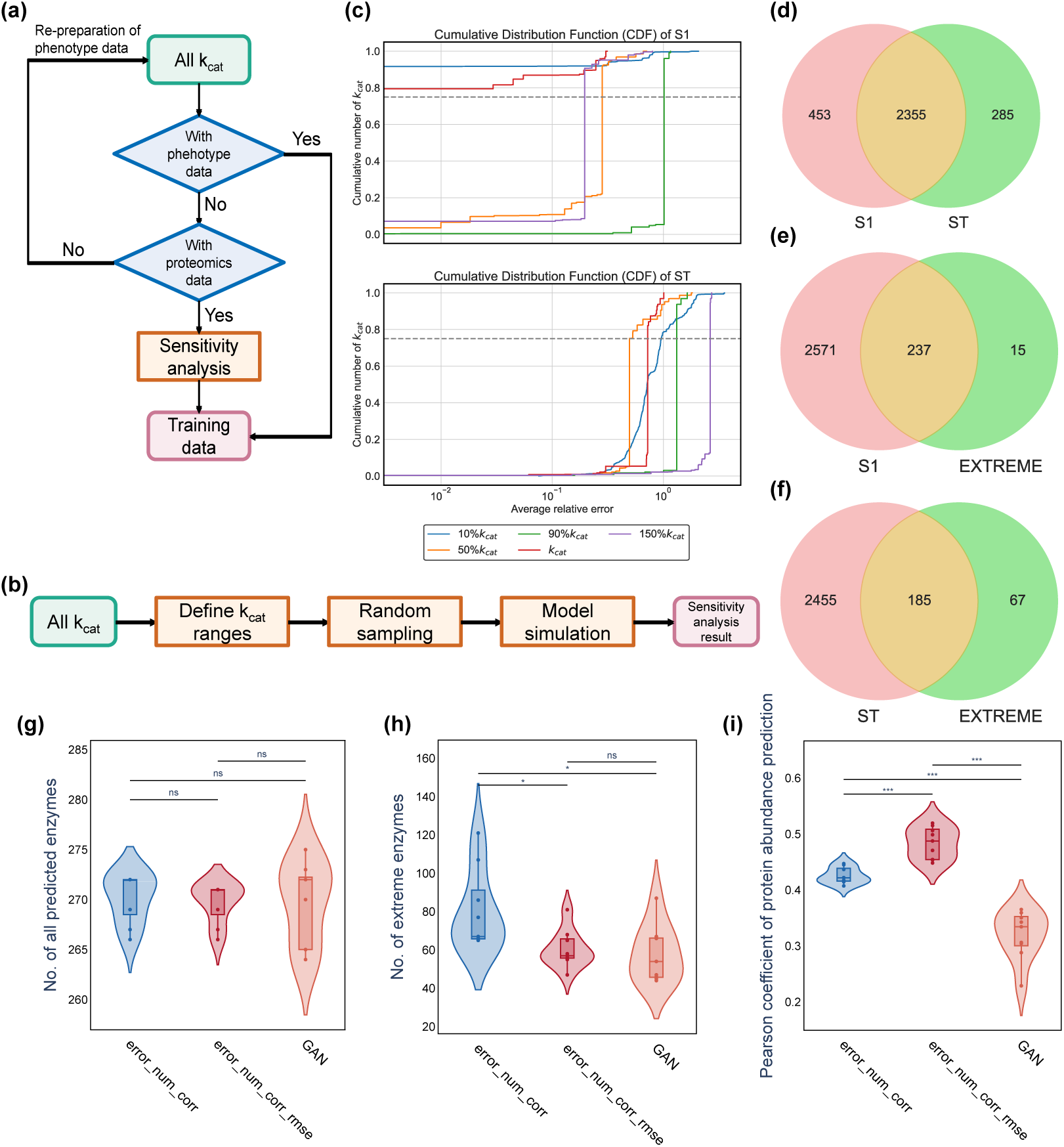
Sensitivity analysis of the *kcat* optimization process and the comparison of the influence of the fitness function. a. Overview of the Generative Adversarial Network (GAN) framework for *kcat* prediction. b. The workflow of the sensitivity analysis. c. Cumulative Distribution Function (CDF) curves for the sensitivity analysis results. S1, first-order sensitivity index; ST, total-order sensitivity index. d. Venn diagram illustrating the intersection of *kcat* values between ST and S1. e. Venn diagram depicting the intersection of *kcat* values between S1 and the set of extreme enzymes.a f. Venn diagram illustrating the intersection of *kcat* values between ST and extreme enzymes. g. Comparison of the total number of predictable enzymes using different fitness function. h. Comparison of the number of extreme enzymes using using different fitness function. i. Comparison of Pearson correlation coefficients of protein abundance predictions between different fitness function. The annotations above the figure represented the t-test results comparing the error_num and error_num_rmse groups, the error_num and GAN groups, and the error_num_rmse and GAN groups, respectively. ns: not significant, *: *P* value <0.05, **: *P* value <0.01, ***: *P* value <0.005.

### Influence of key parameters on EnzymeTuning performances

To effectively improve the protein abundance prediction performance of ecGEMs, three aspects must be taken into consideration: (1) increasing the number of predictable proteins, (2) reducing the number of extreme enzymes, and (3) enhancing prediction accuracy as measured by the PCC. Here, we focused on minimizing extreme enzyme values by defining an appropriate tuning range. Specifically, extreme enzymes were classified into five groups according to the fold change between predicted and measured abundance: >100 or <0.01; >50 or<0.02; >10 or <0.1; >5 or <0.2; and >2 or <0.5. These thresholds corresponded to 198, 258, 369, 424, and 510 extreme enzyme–reaction pairs,

respectively. Strikingly, the group defined by fold changew >50 or <0.02 accounted for over 50% of the total measured abundance of all extreme enzymes, highlighting its priority for optimization. The EnzymeTuning framework was subsequently applied to optimize the *kcat* parameter sets associated with each group (**Figure S2**). Across all groups, the framework achieved robust and steadily converging predictive performance (**Figures S2a–e**). Importantly, the group with fold changes >50 or <0.02 yielded the best training outcome, achieving the highest number of predictable enzymes while maintaining a relatively low number of extreme enzymes (**Figures S2f–h**). Consequently, enzymes with a predicted/measured abundance ratio >50 or <0.02 were designated as the target range for extreme enzyme tuning in subsequent analyses.

In addition, we investigated the impact of the discriminator score used during training on protein abundance prediction accuracy. Initially, we employed a discriminator score that simultaneously maximized the PCC and minimized the fold change between predicted and measured protein abundances (**Eq. 1**). As shown in **Figure S3**, this strategy led to a slight increase in the total number of predictable proteins (**Figure S3a**) and a modest decrease in the number of extreme enzymes (**Figure S3b**), although neither change was statistically significant (*P* value >0.05). By contrast, the Pearson correlation coefficient improved significantly (**Figure S3c**, *P* value <0.05). These results suggested that while using PCC as an evaluation criterion for *kcat* parameter sets enhanced prediction accuracy, directly optimizing fold change limited impact on reducing the number of extreme enzymes. This might be because averaging fold changes in protein abundance predictions diluted the influence of some extreme cases (e.g. fold change >10,000). To address this, we replaced fold change with the number of extreme enzymes as the discriminator score (**Eq. 2**). Furthermore, to ensure generalizability where *kcat* parameters were not pre-adjusted based on growth phenotype prediction performance, we extended the approach to **Eq. 3** by incorporating growth phenotype prediction accuracy (RMSE) alongside protein abundance prediction for all *kcat* parameters.

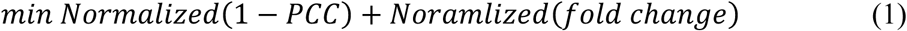

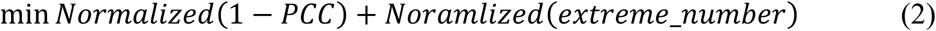

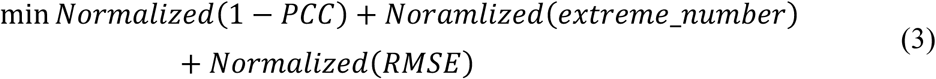

Here, “PCC” referred to the Pearson correlation coefficient for protein abundance prediction, “fold change” was the ratio of predicted to measured protein abundances, and “extreme_number” represented the number of extreme enzymes.

Based on these three discriminator scores, we adjusted the EnzymeTuning framework and compared the resulting protein abundance prediction performance (**Figure 2g-i**). The ecGEMs parameterized with *kcat* sets optimized using **Eq. 2** and **Eq.3** did not exhibit significant changes in the number of predictable proteins; however, both approaches consistently outperformed the **Eq. 1**-based optimization (**Figure 2g**). Across all three methods, simultaneously considering the number of extreme enzymes and growth phenotype prediction (RMSE) led to a significant reduction in the number of extreme enzymes and a marked increase in PCC for protein abundance prediction (**Figures 2h, 2i**). These results highlighted that incorporating growth phenotype prediction accuracy was crucial for constructing realistic *kcat* distributions. Growth phenotype directly reflected the global balance of intracellular metabolic fluxes. Thus, ensuring accurate phenotype predictions indirectly constrained the feasible flux space of the metabolic models, thereby narrowing the solution space for the *kcat* values and guiding them toward biologically realistic enzyme catalytic capabilities. Consequently, **Eq. 3** was selected as the final discriminator score and adopted in the subsequent modeling analyses.

### EnzymeTuning improves the quantitative proteomics prediction

In Step 2, we applied the EnzymeTuning framework to optimize the 237 *kcat* parameter sets identified through sensitivity analysis. The model was trained for 50 iterations, generating a new *kcat* parameter distribution every 10 epochs. Quantitative proteomics datasets under both carbon- and nitrogen-limited conditions, obtained from the literature, were used to validate the generalizability and robustness of the predicted outcomes [20].

As shown in **Figures 3a** and **3b**, the PCC for protein abundance predictions steadily increased, whereas the prediction error consistently decreased across training iterations. These trends indicated that the generator within the GAN framework successfully learned to produce *kcat* distributions yielding progressively more accurate protein abundance predictions. Moreover, the training process exhibited convergence (**Figure 3c**), confirming the reliability and stability of the EnzymeTuning framework. Subsequently, we parameterized the ecGEM with the *kcat* values optimized by EnzymeTuning and compared its predictive performance with the ecGEM parameterized using Bayesian posterior *kcat* values (**Figure 3d–g**). The results revealed a significant increase in PCC for protein abundance predictions (**Figure 3d**) and a marked reduction in the number of extreme enzymes (**Figure 3e**), while maintaining the total number of predicted proteins (**Figure 3f**). These results indicated that EnzymeTuning effectively improved the accuracy of protein abundance predictions. Importantly, similar improvements were also observed under three alternative nitrogen source conditions (glutamate, iso-leucine, and phenylalanine) (**Figure S4**), further validating the generalizability of the framework.

**Figure 3.**
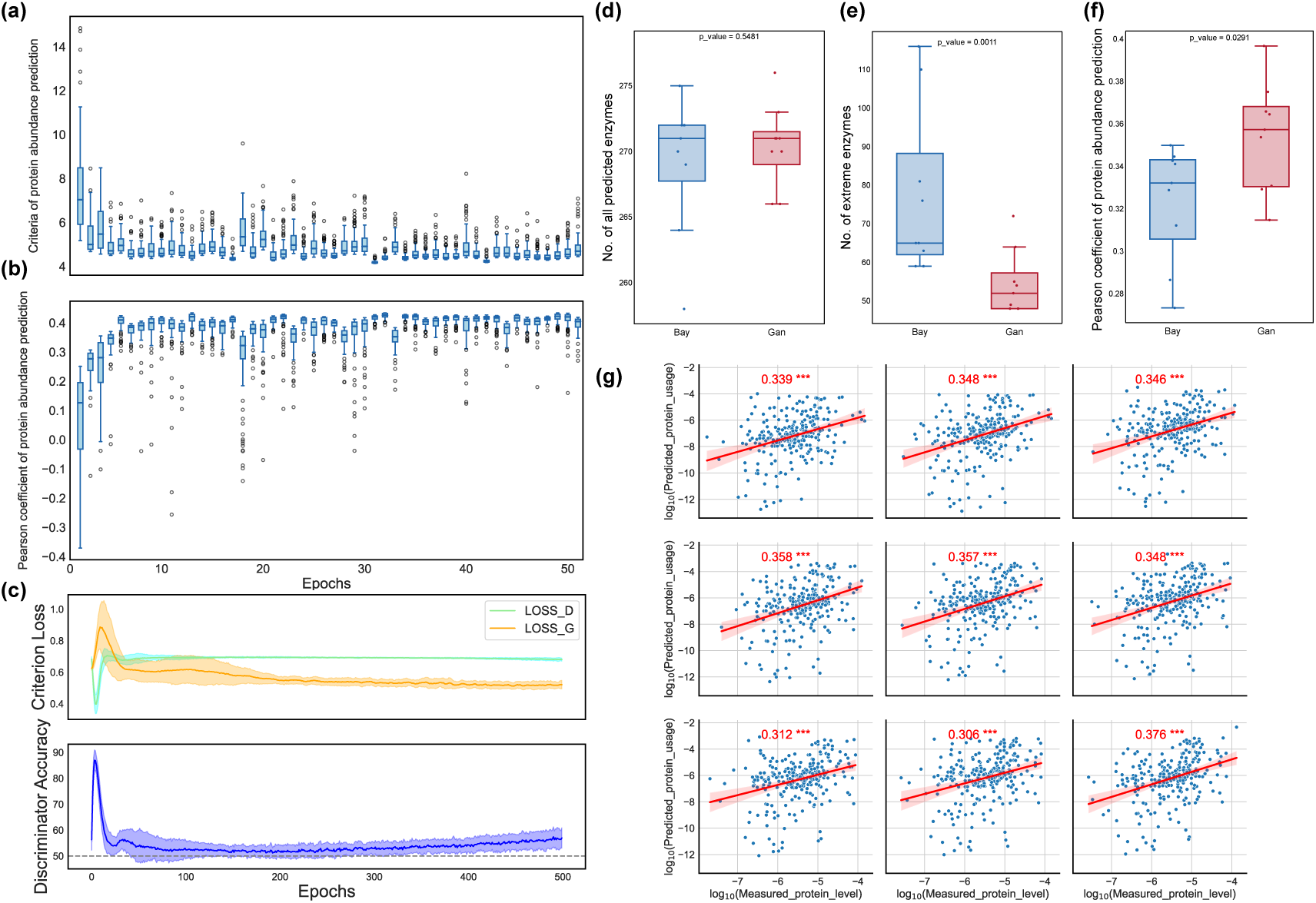
Performance of the GAN approach for protein abundance prediction. a. Changes in Pearson correlation coefficients for protein abundance predictions in the training process. b. Changes in log2(fold change) values during GAN training. c. Discriminator loss (green), generator loss (orange), and discriminator accuracy (blue) over training epochs. d. Comparison of Pearson correlation coefficients of protein abundance predictions between the GAN method and the Bayesian approach. e. Comparison of the number of extreme enzymes using the GAN method and the Bayesian approach. f. Comparison of the total number of predictable enzymes using the GAN method versus the Bayesian approach. g. Pearson correlation coefficients and corresponding *P* values for protein abundance predictions across GAN training. *: *P* value <0.05, **: *P* value <0.01, ***: *P* value <0.005.

Furthermore, we performed KEGG enrichment analysis on the predictable proteins and assessed the protein abundance prediction accuracy for each enriched pathway using PCC (**Figure S5**). As illustrated in **Figure S5a**, proteins with high predictive accuracy (PCC>0.8) were primarily enriched in carbohydrate metabolism pathways—such as glycolysis/gluconeogenesis and fructose and mannose metabolism—as well as amino acid metabolism pathways (e.g., tyrosine and arginine biosynthesis), and nucleotide metabolism. These results suggested that the model effectively captured the correspondence between protein abundance and metabolic flux in these core metabolic modules. Other pathways, such as nitrogen metabolism and glycerolipid metabolism, also demonstrated relatively good predictive accuracy (PCC > 0.5), indicating that the GAN framework exhibited strong sensitivity to central carbon and nitrogen metabolism. By contrast, proteins with low predictive accuracy (PCC < 0.5) were predominantly enriched in pathways such as biosynthesis of secondary metabolites and oxidative phosphorylation (**Figure S5b**). These pathways are typically governed by complex regulatory mechanisms or condition-specific expression patterns, complicating the assignment of consistent *kcat* values across diverse conditions. For instance, enzymes involved in oxidative phosphorylation are highly responsive to mitochondrial activity and respiratory state. However, current ecGEMs do not explicitly incorporate subcellular compartmentalization or dynamic regulation of enzyme activity, potentially contributing to prediction errors. Moreover, broad KEGG categories such as “metabolic pathways” encompass heterogeneous functional modules, further complicating accurate modeling.

To comprehensively evaluate the performance, we compared the protein abundance prediction accuracy between GAN-optimized and Bayesian-optimized ecGEMs across multiple KEGG pathways (**Figure S5c**). In the majority of pathways, GAN-based models achieved higher PCC than Bayesian-based models, demonstrating superior adaptability and generalization. Notably, pathways such as tyrosine metabolism, pyrimidine metabolism, glycerolipid metabolism, and the pentose phosphate pathway exhibited substantial improvements (PCC>0.2) under GAN optimization. These pathways represented central metabolic modules where protein expression closely correlated with flux control, making them tractable for generative modeling approaches. Collectively, these findings supported the conclusion that the EnzymeTuning framework, by learning and generating high-performance *kcat* distributions, effectively captured multi-objective features (e.g., phenotype, protein abundance, and extreme enzyme regulation), thereby yielding ecGEMs with enhanced physiological realism.

### Estimation of protein synthesis rates (*vsyn*) for unused proteins inferred by EnzymeTuning

Despite optimization with EnzymeTuning, the ecGEM did not show a significant increase in the number of predictable proteins. In the *S. cerevisiae* ecGEM, 1,151 metabolic enzymes are represented, while only 200–300 enzymes had their abundances predicted, leaving many enzymes unaccounted for. This limitation might arise because most metabolic reactions in the genome-scale network lack constraints, rendering them inactive (flux = 0). To address this, we sought to broaden the predictive scope by constraining enzyme abundances using predicting protein synthesis rates (*vsyn*), which were derived from literature-reported protein degradation constants (*kdeg*) (**Figure 4a**) [7, 21].

**Figure 4.**
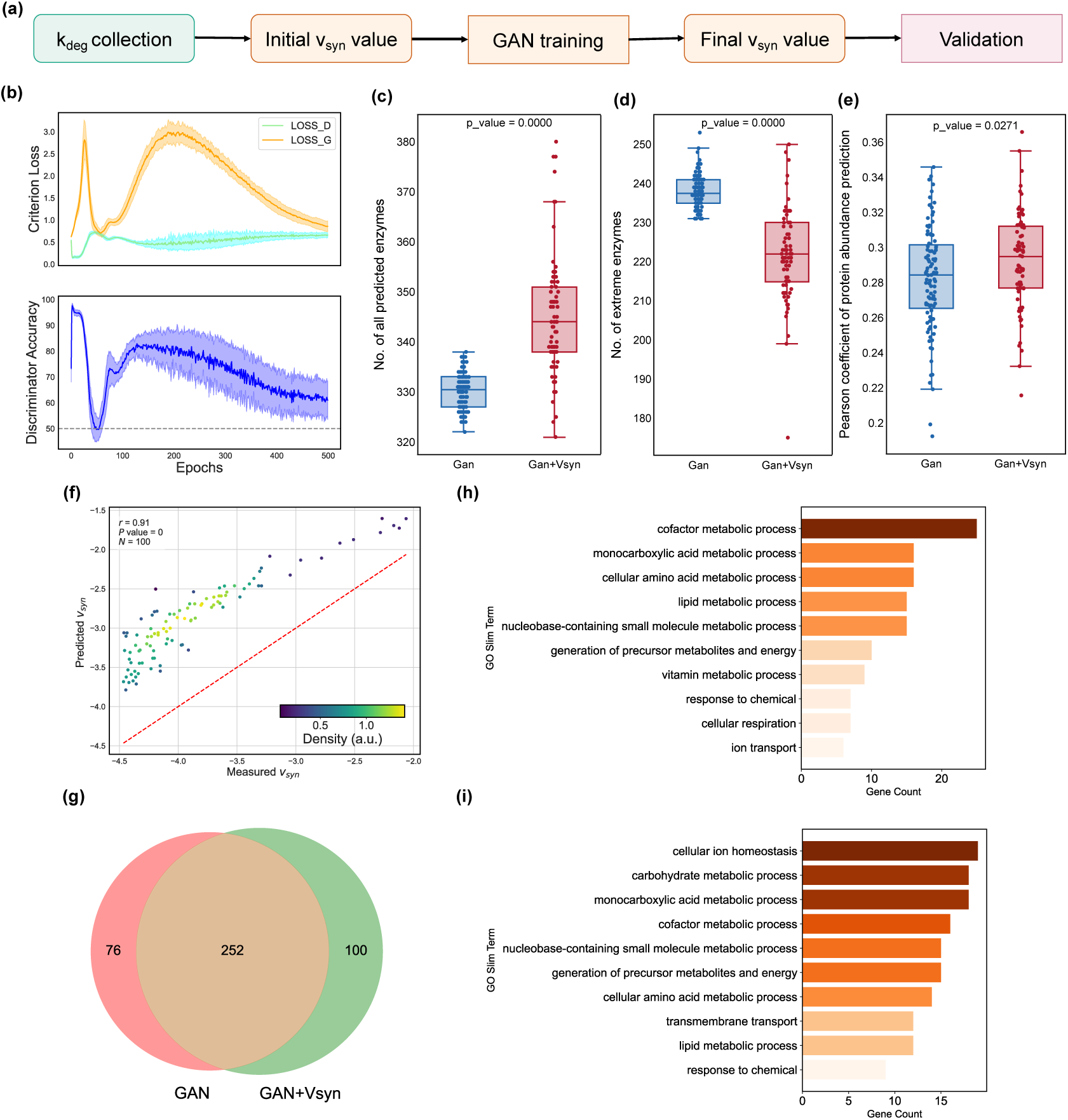
Performance of the GAN approach for protein synthetic rate prediction. a. The workflow of the GAN approach for protein synthetic rate prediction. b. Discriminator loss (green), generator loss (orange), and discriminator accuracy (blue) over training epochs. c. Comparison of the total number of predictable enzymes using the GAN method versus the GAN approach for protein synthetic rate. d. Comparison of the number of extreme enzymes using the GAN method and the GAN approach for protein synthetic rate. e. Comparison of Pearson correlation coefficients of protein abundance predictions between the GAN method and GAN approach for protein synthetic rate. f. Correlation between GAN-predicted *vsyn* values and *vsyn* values calculated via **Eq.5**. *P* value for the Pearson correlation coefficients was calculated using Student’s t-test. g. Venn diagram representing the overlap between predicted proteins between GAN method and GAN method with protein synthetic rate (GAN+*Vsyn*). h and i. the GO enrichment results of proteins uniquely predicted by GAN method (h) and GAN- *Vsyn* method (i).

Using protein mass conservation relationships and *kdeg* values, we converted degradation rates into corresponding protein synthesis rates (*vsyn*) (**Eq. 5–8**). Initially, we constrained the ecGEM with all available *vsyn* values; however, this over-constraining markedly reduced predictive performance. Consequently, we sought an optimal range of *vsyn* constraints that maximized improvements in protein abundance prediction. To this end, we identified proteins with predicted abundances of zero, ranked the experimentally measured protein abundances in descending order, and selected the top 10, 25, 50, 100, 200, 300, and 400 proteins as candidate datasets for *vsyn* optimization. The comparative analyses (**Figure S6**) revealed that constraining the top 100 *vsyn* values provided the best trade-off between protein abundance prediction and phenotype prediction accuracy. These optimized *vsyn* values were subsequently used to define the upper and lower bounds of enzyme exchange reactions.

As shown in **Figure 4b**, the EnzymeTuning framework exhibited stable convergence during *vsyn* optimization, confirming the stability of the learned *kcat* distributions. To evaluate the effect of introducing additional constraints, we compared two ecGEM variants under batch cultivation: (1) parameterized solely with GAN- optimized *kcat* values, and (2) further constrained with *vsyn*-derived protein synthesis bounds. The *vsyn*-constrained ecGEM demonstrated superior predictive performance with the number of predictable enzymes increasing to ∼380 and a marked reduction in the number of extreme enzyme values (**Figures 4c–e**). Collectively, these improvements highlighted the potential of *vsyn* constraints to substantially expand the predictive scope and accuracy of ecGEMs.

Comparison of protein synthesis rates (*vsyn*) estimated by ecGEM and those derived from protein degradation constants (*kdeg*) revealed that the ecGEM-predicted *vsyn* values were generally higher (**Figure 4f**). This discrepancy reflected that ecGEM simulations focused primarily on protein synthesis without fully accounting for protein degradation. The *vsyn* represented the total rather than net synthesis rates, thus potentially leading to overestimation. Nevertheless, the two approaches were strongly correlated (*r* = 0.91) indicating that ecGEM reliably captured the relative variations in synthesis rates despite absolute offsets. This was consistent with previous findings that protein synthesis rates were major determinants of protein abundance [22]. **Figure S7** showed strong agreement between the predicted and measured protein synthesis rates. The density distribution (**Figure S7a**) demonstrated that the model accurately reproduced the overall *vsyn*. Residual analysis (**Figure S7b**) revealed small and unbiased errors across most of the range, with slightly elevated variance at high synthesis rates. **Figure S7c** further confirmed a high correlation between predicted and measured values (R² = 0.90).

To identify which proteins benefited from the incorporation of *vsyn* constraints, we compared the predicted protein abundances between the *vsyn*-constrained GAN-ecGEM and the original GAN-ecGEM. Gene Ontology (GO) enrichment analysis was then performed for proteins uniquely predictable by each model (**Figure 4h-i**). Proteins predicted exclusively by the GAN-ecGEM were primarily enriched in biological processes associated with core metabolism and cellular maintenance functions, such as "cofactor metabolic process", and "cellular amino acid metabolic process" (**Figure 4h**). Interestingly, several peripheral or conditional-specific categories, such as “conjugation”, “meiotic cell cycle”, and “regulation of cell cycle”, were also enriched. The enrichment of these GO terms suggested that the GAN-ecGEM retained flexibility in capturing non-canonical expression patterns.

By contrast, proteins recovered only after introducing *vsyn* constraints were enriched in "cellular ion homeostasis" (**Figure 4i)**, a category typically characterized by low expression, high variability, and strong post-transcriptional regulation [23]. This category typically comprised transporters such as H⁺/Na⁺ pumps and metal ion transporters, whose abundances were sensitive to environmental fluctuations and translational regulation [24]. Such proteins often exhibited inherently low expression and high variation, making them challenging to predict consistently. The incorporation of *vsyn* constraints improved the model’s ability to capture these "functionally peripheral" proteins, enabling their accurate prediction. These fundings further supported the effectiveness of incorporating the *vsyn* constraints to expand the predictive capacity of ecGEMs.

### Automatic reconstruction high-quality ecGEMs for various organisms using EnzymeTuning

The EnzymeTuning framework successfully optimized the *kcat* parameter set in Crabtree-positive *Saccharomyces cerevisiae*. We next assessed its applicability for enzyme kinetic parameter tuning in three Crabtree-negative yeasts (*Yarrowia lipolytica*, *Kluyveromyces marxianus*, and *Kluyveromyces lactis*) to evaluate its generalizability across species with distinct metabolic strategies. These species were selected to highlight the broad utility of EnzymeTuning. Enzyme-constrained models for each microorganism have been constructed, providing the foundational for ecGEM modeling. Parameter optimization was then conducted following the EnzymeTuning workflow, using published experimental data for growth phenotypes and quantitative proteomics [25]. As shown in **Figure 5a-c**, EnzymeTuning consistently improved predictive performance across all three species. In *K. lactis,* the number of predictable proteins significantly increased, with the median rising from 240 to 247 (*P* value <0.05; **Figure 5a**). **Figure 5b** demonstrated a clear reduction in the number of extreme enzymes in all three yeasts. For example, in *Y. lipolytica*, the number of extreme enzymes dropped from ∼65 to ∼40 (*P* value <0.05), indicating that EnzymeTuning alleviated reliance on outlier enzyme parameters. Correspondingly, prediction accuracy improved in all three yeasts, with PCC values increasing from ∼0.33 to ∼0.37 in *K. marxianus* (*P* value <0.05), from ∼0.21 to ∼0.29 in *K. lactis*, reflecting enhanced proteome-level performance (**Figure 5c**). Collectively, these results further validate the robustness of the EnzymeTuning in improving the predictive capability and demonstrate its applicability to non-model organisms, supporting the metabolic optimization in industrial strains.

**Figure 5.**
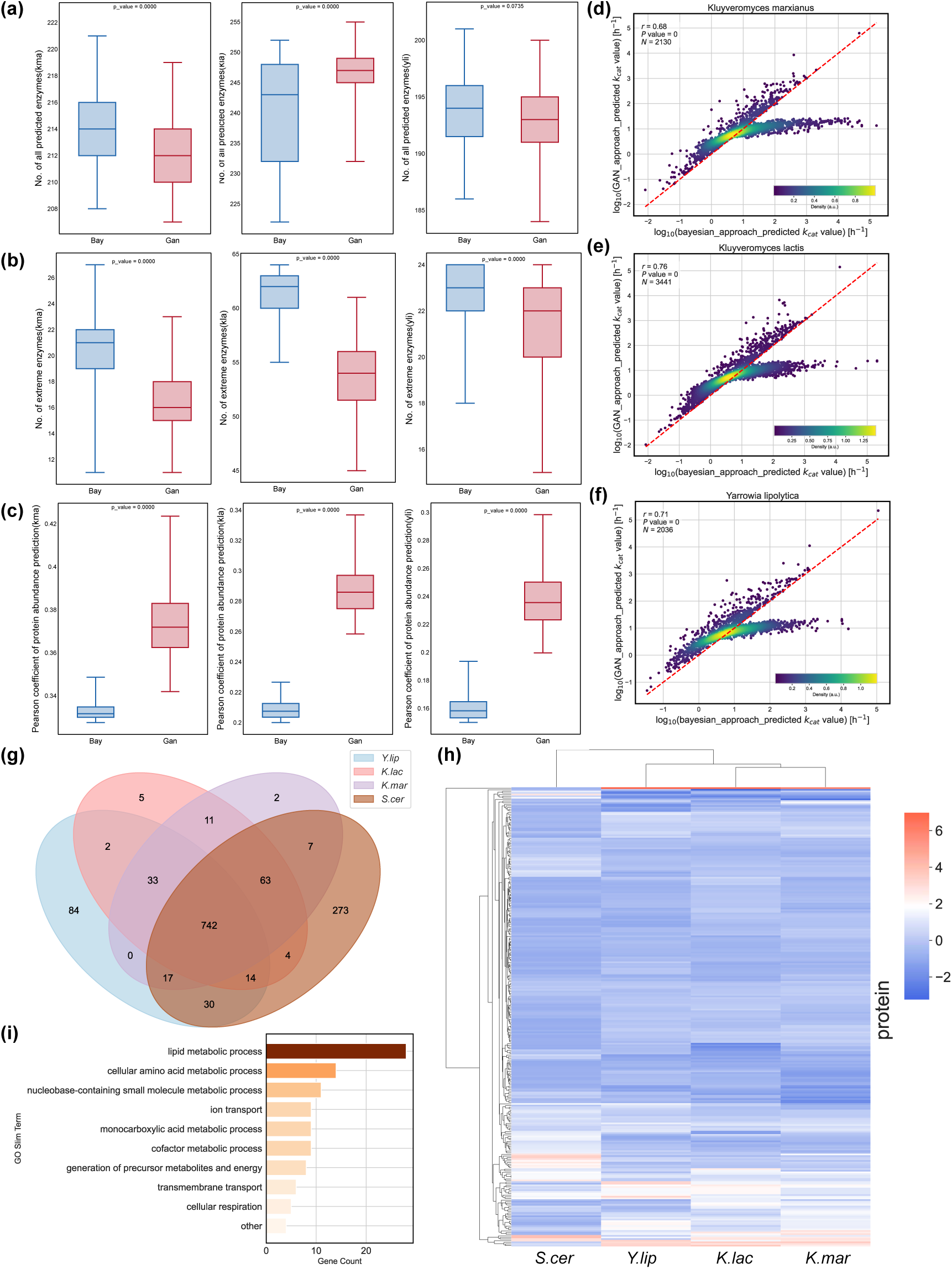
Performance of the GAN approach in other yeasts and comparative Analysis of *kcat* distributions across crabtree-positive and crabtree-negative yeasts. a. Comparison of the total number of predictable enzymes using the GAN method versus the Bayesian approach in *K. marxianus*, *Y. lipolytica*, and *K. lactis*. b. Comparison of the number of extreme enzymes using the GAN method and the Bayesian approach in *K. marxianus*, *Y. lipolytica*, and *K. lactis*. c. Comparison of Pearson correlation coefficients of protein abundance predictions between the GAN method and the Bayesian approach in *K. marxianus*, *Y. lipolytica*, and *K. lactis*. d–f. Density scatter plots comparing predicted *kcat* values between the GAN and Bayesian approaches for the three crabtree-negative yeasts: *K. marxianus* (d), *K. lactis* (e), and *Y. lipolytica* (f). g. Venn diagram showing the overlap of genes used in the models across the four yeasts (*S. cerevisiae*, *K. marxianus*, *K. lactis*, and *Y. lipolytica*). h. Heatmap of *kcat* values for the shared genes across the four species. i. GO term enrichment of genes with significantly different *kcat* values between crabtree- negative (non-*S. cerevisiae*) and crabtree-positive (*S. cerevisiae*) species, showing the top 10 enriched GO-slim terms.

Subsequently, we compared *kcat* distribution optimized by GAN and Bayesian methods across the three yeast species (**Figure 5d–f**). Density plots revealed overall positive correlations between the two approaches. However, *kcat* values predicted by

GAN tended to be lower in specfic regions. This discrepancy arised because GANs were trained to capture the full distribution, including the tails of real distributions (e.g., lower-value regions). In contrast, the Bayesian inference typically yielded smoother estimates that might overestimate *kcat* values in sparse regions.

To further investigate the enzymatic catalytic capacities, we analyzed 742 proteins shared a mong the three Crabtree-negative yeasts and the Crabtree-positive *S. cerevisiae* (**Figure 5g**). Their corresponding *kcat* values were shown in a heatmap (**Figure 5h**). Correlation analysis revealed strong consistency among the three Crabtree-negative species (PCC *r*>0.9), indicating high conservation of enzymatic efficiency within this group, consistent with their predominantly reliance on respiratory metabolism. This observation aligned well with previous reports showing that Crabtree-negative yeasts exhibited elevated mitochondrial respiratory capacity and oxidative phosphorylation [26].

In contrast, pronounced differences were observed when comparing the Crabtree-negative species with *S. cerevisiae*. The GO enrichment analysis of proteins with differential *kcat* values (**Figure 5i**) revealed enrichment in the functional categories, including lipid metabolism, cellular amino acid metabolism, nucleobase-containing small molecule metabolism, ion transport, and cofactor metabolism. A comparative analysis of *kcat* distributions among proteins co-expressed by three Crabtree-negative yeasts (*K. lactis*, *K. marxianus*, and *Y. lipolytica*) and the Crabtree-positive yeast *S. cerevisiae* was investigated (**Figure 5h, i**). On average, Crabtree-negative yeasts exhibited higher catalytic efficiencies for enzymes involved in amino acid metabolism but lower efficiencies for lipid metabolism enzymes compared with *S. cerevisiae*. These trends likely reflected fundamental differences in metabolic strategies, resource allocation, and physiological adaptations. In Crabtree-negative yeasts, the *kcat* values observed for amino acid metabolic enzymes might support their strong reliance on aerobic respiration, enabling higher growth rates and enhanced protein synthesis [27]. Enhanced mitochondrial function and nitrogen utilization capacity in these yeasts necessitated efficient amino acid biosynthesis to satisfy the cellular demand for nitrogen-containing intermediates. Conversely, during fermentative metabolism, *S. cerevisiae* constantly remodeled its membrane systems (e.g., plasma membrane, endoplasmic reticulum) to counter fluctuations in intracellular pH and osmotic pressure [28]. This dependence on lipid remodeling might impose higher catalytic efficiency requirements on lipid metabolic enzymes. Overall, these findings suggested that Crabtree-negative yeasts prioritized catalytic efficiency toward amino acid metabolism, reflecting a strategic trade-off in enzyme resource allocation to optimize energy and nutrient utilization under distinct metabolic regimes.

### Estimation of enzyme turnover numbers (*kcat*) across different carbon-nitrogen conditions

To further investigate the regulatory mechanisms underlying enzyme catalytic capacity under varying carbon-to-nitrogen (C/N) conditions, we leveraged literature-derived proteomics datasets and trained GAN models at four distinct C/N ratios (3, 30, 50, 115) to generate condition-specific *kcat* distributions [25]. In this classification, CN3 was designated as carbon-limited conditions (C-lim), whereas CN30, CN50 and CN115 were categorized as nitrogen-limited conditions (N-lim). For subsequent comparative analyses, CN3 and CN115 were selected as representative C-lim and N-lim conditions, respectively. To evaluate whether the overall *kcat* distributions differed significantly across nutrient conditions, we employed the Kolmogorov–Smirnov (KS) test, a non-parametric statistical method for comparing continuous distributions between two groups [6]. No significant global differences were detected, as confirmed by the KS tests between C-lim and N-lim states (KS-stat = 0.076, p = 0.98). Given that *in vitro* estimates (i.e., *kcat*) of the maximal enzyme activities predicted often failed to capture the actual intracellular utilization states of enzymes, we introduced a relative enzyme usage metric [29]. Specifically, for each enzyme under a given condition, we computed the ratio of its condition-specific predicted *kcat* to the maximum *kcat* value (𝑘^𝑚𝑎𝑥^) reported in the previous studies, thereby obtaining the normalized ^𝑘𝑐𝑎𝑡^/ _𝑚𝑎𝑥_ ratio as a proxy for relative enzyme usage.

Partial least squares discriminant analysis (PLS-DA) based on the relative enzyme usage metric was conducted to identify proteins contributing most to group differentiation (**Figure 6a**) [30]. Proteins were ranked by their Variable Importance in Projection (VIP) scores, and the top 20 proteins were selected for functional analysis (**Figure 6b**). Functional annotation revealed that these proteins primarily participate in central carbon metabolism, amino acid metabolism, lipid metabolism, and cell wall biogenesis. Specifically, several enzymes were associated with lactate metabolism (e.g., CYB2, DLD1, DLD2), highlighting the potential metabolic adaptation to fluctuating redox and energy demands. Key enzymes in branched-chain amino acid (BCAA) and aromatic amino acid biosynthesis (e.g., LEU1, LEU9, and ARO3) were also identified, suggesting reprogramming of nitrogen metabolism under different C/N regimes. Interestingly, multiple proteins (e.g., FKS1, FKS2, KRE6, SKN1) were associated with β-glucan synthesis and fungal-type cell wall organization, suggesting that structural remodeling of the cell envelope represented a major adaptive strategy under nutrient variation.

**Figure 6.**
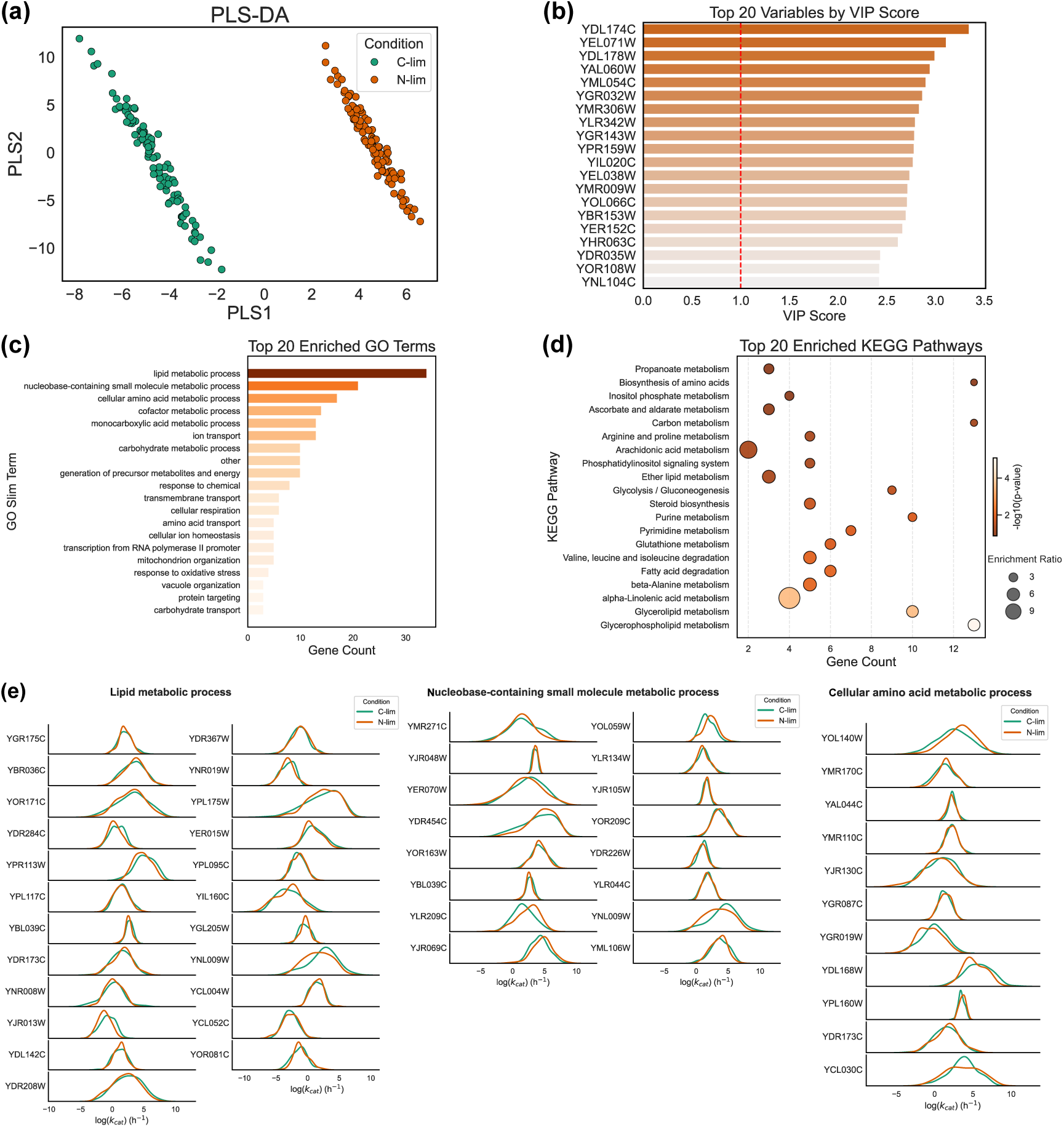
The *kcat* distribution and functional enrichment analysis of differential proteins under various C/N conditions. a. Partial Least Squares Discriminant Analysis (PLS-DA) of *kcat* values under C-lim and N-lim conditions. Each point represents a sample, colored by the corresponding nutrient condition (C-lim or N-lim). b. Variable Importance in Projection (VIP) scores of *kcat* parameters derived from the PLS-DA model. The top 20 enzymes with the highest contribution to group discrimination were shown. c. Gene Ontology (GO) enrichment analysis of differential proteins. d. KEGG pathway enrichment analysis of differential proteins. e. The *kcat* distribution of differential proteins grouped by GO terms.

As shown in **Figure 6c-e**, GO and KEGG enrichment analysis were performed to further characterize the condition-sensitive enzyme usage[31]. Consistent with the PLS-DA results, the GO analysis revealed significant enrichment in lipid metabolism, nucleobase-containing small molecule metabolism, amino acid metabolism, cofactor metabolism, and mitochondrial organization (**Figure 6c**). Notably, the metabolic processes associated with energy generation, respiratory chain activity, as well as fatty acid metabolism were significantly enriched, underscoring the reprogramming of both energy supply and membrane-related pathways under varying C/N conditions. Consistently, KEGG pathway analysis highlighted propanoate metabolism, glycolysis, and aromatic amino acid biosynthesis as key pathways responsive to carbon-nitrogen balance (**Figure 6d**). Notably, aromatic amino acid biosynthesis played a pivotal role in maintaining protein synthesis capacity and energy homeostasis under nitrogen-limited conditions [32]. These results suggested that C/N-driven adjustments in catalytic efficiency were not restricted to primary carbon and nitrogen metabolism but extend to a broad network of energy, lipid, and cofactor pathways. This reflected a coordinated metabolic adaptation strategy to optimize growth and survival under fluctuating nutrient environments.

Further, we analyzed the density distributions of ^𝑘𝑐𝑎𝑡^/ _𝑚𝑎𝑥_ ratios for representative enzymes from three key functional categories under different conditions (**Figure 6e**). Under N-lim conditions, the lipid metabolism module showed significantly reduced relative usage. These proteins were primarily involved in glycerophospholipid/GPI-anchor and sphingolipid pathways, (such as (GPI)-anchor initiation (PIS1, GPI3, GPI14), ergosterol and sterol-ester formation (ERG1, ARE2) and triglyceride remodeling and phosphatidate turnover (LRO1, DPP1)). This finding indicated that, under nitrogen limitation, cells actively suppressed enzymatic activities within energetically costly lipid metabolism pathways to maintain energy balance. Such regulatory patterns were consistent with the mechanisms of post-transcriptional and enzyme-level regulation that suppress lipid biosynthetic processes under nutrient stress conditions [33]. Notably, Xia et al. reported a similar trend under carbon limitation, where increasing specific growth rates also were accompanied by reduced proteome allocation to lipid biosynthesis, indicating a general regulatory principle in *S. cerevisiae* across distinct elemental limitations [34].

Additionally, enzymes involved in amino acid metabolism also displayed N-lim conditions, whereas rnithine acetyltransferase (ARG8) and aldehyde dehydrogenase (ALD2) displayed increased relative usage. ARG8 facilitated nitrogen recycling by channeling nitrogen into the glutamate–2-oxoglutarate pool, consistent with the role of the Arg/Orn cycle as a mobilizable nitrogen reservoir under scarcity [35]. ALD2 was essential for maintaining CoA supply and redox balance under nutrient stress [36]. Conversely, enzymes associated with glycolysis, glycerol metabolism, and nucleotide catabolism showed increased utilization under N-lim conditions, including glycerol-3- phosphate dehydrogenase (GPD2), and inosine triphosphate pyrophosphatase (ITPA). Together, these patterns indicated a coordinated metabolic strategy in which *S. cerevisiae* suppressed energetically costly biosynthetic routes while up-regulating pathways that support nitrogen recycling, redox homeostasis, and precursor metabolite maintenance. Such adaptations were consistent with previously described metabolic remodeling mechanisms under nitrogen stress [25].

## Discussion

In this study, we developed a generative adversarial network (GAN)-based framework, EnzymeTuning, to optimize *kcat* parameter sets in enzyme-constrained genome-scale metabolic models (ecGEMs). Compared with Bayesian posterior sampling approaches, the EnzymeTuning framework significantly improved the accuracy of protein abundance predictions while maintaining robust phenotype predictions [11]. This demonstrated the potential of GANs to capture complex, nonlinear *kcat* distributions and to simultaneously optimize multiple biological objectives [37]. Unlike Bayesian inference, which yields maximum a posteriori estimates of *kcat* values based on prior assumptions, the GAN-based approach can learn parameter distributions driven jointly by global constraints (e.g., phenotype fidelity) and local evaluation (e.g., reduction of extreme enzyme burden, correlation with proteomics). We demonstrated the generalizability and robustness of the framework by extending its application from *S. cerevisiae* to three Crabtree-negative yeasts (*K. lactis*, *K. marxianus*, and *Y. lipolytica*).

Nevertheless, extreme enzyme predictions were still concentrated in complex regulatory pathways such as oxidative phosphorylation. These cases may reflect inherent limitations of the framework, including the lack of cellular compartmentalized constraints, isoenzyme-level differentiation, and post-translational regulatory information [38, 39]. In addition, training GANs for high-dimensional enzyme networks remains computationally intensive, and the interpretability of the generated parameter distributions is limited. To alleviate bottlenecks in protein predictability, we incorporated the protein synthesis rate (*vsyn*), which was inferred from protein degradation coefficients (*kdeg*), as auxiliary constraints, increasing the predictable enzyme coverage in the *S. cerevisiae* ecGEM from ∼290 to ∼380 (**Figure 4c**) [21, 40]. To further explore the physiological mechanisms underlying condition-specific enzyme utilization, we applied EnzymeTuning to proteomics datasets collected under four distinct carbon-to-nitrogen (C/N) ratios (3, 30, 50, and 115). By analyzing *in vivo* enzyme catalytic efficiency through the ^𝑘𝑐𝑎𝑡^/ _𝑚𝑎𝑥_ ratio, we revealed adaptive enzyme usage patterns consistent with nutrient-dependent resource allocation strategies. The incorporation of regulatory layers (such as phosphor-proteomics, compartment-specific kinetics, or environmental parameters like pH, osmotic stress, and oxygen availability) may further enhance the model fidelity. The framework can be extended to additional perturbation conditions to capture broader patterns of enzymatic resource allocation and metabolic plasticity. Moreover, integrating EnzymeTuning into high-throughput design-build-test-learn (DBTL) cycles offers the potential to accelerate industrial strain development. Together, these developments point toward a comprehensive and dynamic framework for quantitative, context-specific metabolic modelling.

## Methods

### Dataset resources

The genome-scale metabolic models (GEMs) for *S. cerevisiae*, *Y. lipolytica*, *K. lactis* and *K. marxianus* used in this study were automatically reconstructed from a yeast/fungi pan-GEM using the established modeling framework developed by Lu et al. [41]. Experimental growth phenotypes, exchange flux rates, and the absolute quantitative proteomics data for *S. cerevisiae* under 23 steady-state chemostat cultures, including 9 carbon-limited and 14 nitrogen-limited conditions, were collected from the literatures [20, 34]. The experimental data for other three yeasts were collected from the literatures [42, 43]. Condition-specific proteomic data for *S. cerevisiae* under varying C/N ratios were also obtained from the literature [25]. In this study, all cultures were maintained at a dilution rate of 0.2 h^-1^, and the nitrogen availability was progressively reduced by decreasing ammonium concentration while keeping glucose constant. A C/N ratio of 3 corresponds to a carbon-limited condition, while C/N ≥ 5 reflects a nitrogen-limited state [25]. All these data are available in the GitHub repository (https://github.com/hongzhonglu/EnzymeTuning).

### Sensitivity analysis

In this study, the enzyme-constrained genome-scale metabolic model (ecGEM) of *S. cerevisiae* comprised 4,354 *kcat* parameters. While, Li et al. previously optimized these parameters for phenotype predictions[11], we aimed to improve protein abundance prediction accuracy while maintaining phenotype fidelity. Thus, we introduced a sensitivity analysis-based refinement strategy as an optional step prior to global optimization. This step allowed for targeted improvement of protein abundance prediction when phenotype optimization was not required. We evaluated protein abundance prediction performance from two complementary aspects: (1) the correlation between predicted and measured abundance, assessed by the Pearson correlation coefficient; and (2) the physiological plausibility of enzyme allocation, quantified by the number of extreme enzymes. Here, the extreme enzymes were defined as those with predicted/measured protein abundance ratios outside a predefined range. Enzymes with a fold change greater than 50 or less than 0.02 were considered extreme, with the thresholds determined empirically and described earlier in the manuscript. We employed SALib (Sensitivity Analysis Library in Python) to systematically identify critical *kcat* parameters [19]. Each *kcat* value was perturbed across five intervals (10%- 30%, 50%-70%, 90%-110%, 110%-130%, and 150%-170% of their nominal values). Sampling was performed via the Sobol method to ensure quasi-random coverage of the parameter space. The prediction under these perturbations yielded growth rates and exchange fluxes, with the accuracy quantified by the root mean square error (RMSE) between predicted and experimental datasets (**Eq. 4**).

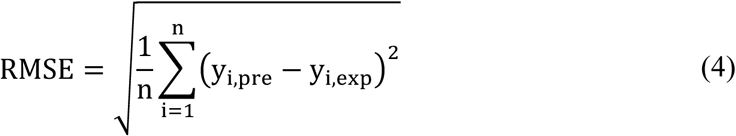

Here, 𝑦_𝑖,𝑝𝑟𝑒_ represents the predicted exchange rate, while 𝑦_𝑖,𝑒𝑥𝑝_ is the experimentally measured value. To access parameter sensitivity, the first-order (S1) and the total-order (ST) indices were computed. Through the S1 and ST results across five perturbation ranges, a pre-optimized *kcat* dataset was generated.

ecGEM reconstruction based on Bayesian methods A sequential Monte-Carlo-based approximate Bayesian computation (SMC-ABC) approach was employed for sampling *kcat* values[44]. The posterior mean of the sampled *kcat* values was then used to parameterize the ecGEM, resulting in the posterior-mean-ecGEM. Further methodological details were available in the work of Li et al. [11].

### Construction of EnzymeTuning

Generative adversarial network (GAN)-based optimization named EnzymeTuning was implemented in three steps to refine enzyme turnover numbers (*kcat*) and enhance predictive performance of ecGEMs (**Figure 1**). The process was initiated with an input set of *kcat* parameters obtained from a Bayesian inference framework [11]. In the first step (**Step 1: Preparation of training data**), each parameter set was systematically evaluated for its ability to simultaneously reproduce two cellular features: (1) growth phenotypes and (2) quantitative protein abundance profiles. Those parameter sets that yielded accurate growth rate predictions and exhibited high correlations with measured protein abundance predictions were classified as high-performance, whereas those failing to meet either criterion were designated as low-performance. When sensitivity analysis was applied prior to global optimization, only protein abundance prediction accuracy was considered. In this context, the high-performance sets were defined solely by their agreement with measured protein abundance profiles. This classification produced a labeled dataset with clear performance-based categories, which was subsequently used for supervised generative modeling.

In the second step, using the labeled dataset, we trained a conditional generative adversarial network (cGAN) (**Step 2: Training the adversarial networks**)[18]. The cGAN consisted of two fully connected neural networks: a generator and a discriminator, which conditioned on the labels during the training. Conditioning enabled the generator to capture the distribution of high-performance parameter sets. In this step, the aim was to produce novel, high-quality *kcat* datasets that emulate the distributions of biologically meaningful parameter sets. All implementations were carried out using Keras with TensorFlow-GPU backend. The discriminator consisted of three dense layers of increasing complexity: the first layer contained 32 units, 64 units, and 128 units, each followed by a Dropout layer (rate = 0.5). The generator network comprised three dense layers with progressive depth and regularization: 128 units, 256 units, and 512 units, each followed by Batch Normalization and Dropout (0.5). Both networks were trained using binary cross-entropy loss and the Adam optimizer with a learning rate of 0.0001. After each training iteration, the generator produces new *kcat* sets, which were integrated into the ecGEM and re-evaluated for its predictions of growth phenotypes and protein abundance. Only the top 100 *kcat* datasets were retained for subsequent iterations over 50 training rounds, progressively converging toward high-performing *kcat* parameter spaces.

The final generator was used to produce *kcat* datasets for performance testing under multiple environmental conditions (**Step 3: Validation**). Model evaluation was conducted in predicting protein abundances of *S. cerevisiae* under carbon-limited conditions to confirm that the generated parameters reproduced the high correlations with measured protein abundances. To further assess the robustness, the generated *kcat* parameter sets were validated under nitrogen limitation and 3 other nitrogen sources (glutamate, iso-leucine, and phenylalanine).

### Vsyn estimation based on the degradation rate

Assuming steady-state intracellular protein levels, we applied the principle of protein mass conservation to estimate the protein synthesis rate (*vsyn*) [45]. According to this principle, for each enzyme, the rate of protein synthesis must balance the sum of degradation and dilution rates (**Eq. 5-8**). The initial *vsyn* values were derived from literature-reported protein degradation constants (*kdeg*) [21, 40]. To account for methodological variations in *kdeg* determinations across different studies, we calculated two boundary estimates (*vsyn,min*,*vsyn,max*) using **Eq. 8**. For ecGEM model construction, the top 100 metabolic enzymes were identified through the following criteria: (1) descending ranking by experimentally measured protein abundance, and (2) predicted protein abundance equal to zero. Specifically, enzymes satisfying both criteria were further ranked by their experimental abundance, and the top 100 were retained. The bounds of protein exchange reactions in the ecGEM were parameterized using these *vsyn* estimates. Finally, the EnzymeTuning framework was applied to systematically calibrate the protein synthesis rates, with the optimized values being integrated into the ecGEM.

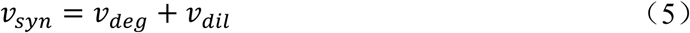

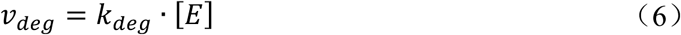

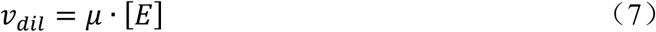

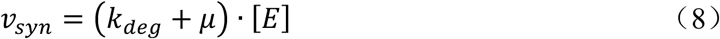

Here, **Eq.5** defined the total protein synthesis rate (*vsyn*). Each protein could be degraded (*vdeg*, **Eq.6**) and be diluted by growth rate (*vdil*, **Eq.7**). **Eq.6** descried the protein degradation as a first-order process, with *kdeg* denoting the protein-specific degradation constant and [𝐸] representing the intracellular protein concentration. **Eq.7** captured the dilution effect of biomass growth, where 𝜇 was the specific growth rate. By combining **Eq.5** to **Eq.7**, **Eq.8** provided a unified expression for the protein synthesis rate which integrated both degradation and dilution effects.

Computational Environment

The EnzymeTuning framework was implemented in Python, with enzyme constrained model reconstruction, training, and evaluation. The complete execution of EnzymeTuning framework required approximately one day to complete. The computations in this paper were run on the Siyuan-1 cluster supported by the Center for High Performance Computing at Shanghai Jiao Tong University.

## Data availability

The data in this study are available in the GitHub repository (https://github.com/hongzhonglu/EnzymeTuning).

## Code availability

We provide all codes and detailed instruction in the GitHub repository (https://github.com/hongzhonglu/EnzymeTuning).

## Acknowledgement

This work was financially supported by Shanghai Municipal Science and Technology Major Project, the National Key R&D Program of China (2022YFA0913000, 2020YFA0908300), Natural Science Foundation of Shanghai (25ZR1402110), grants 22208211 and 22378263 from the National Natural Science Foundation of China.

---

**Supplemental Figure 1.**
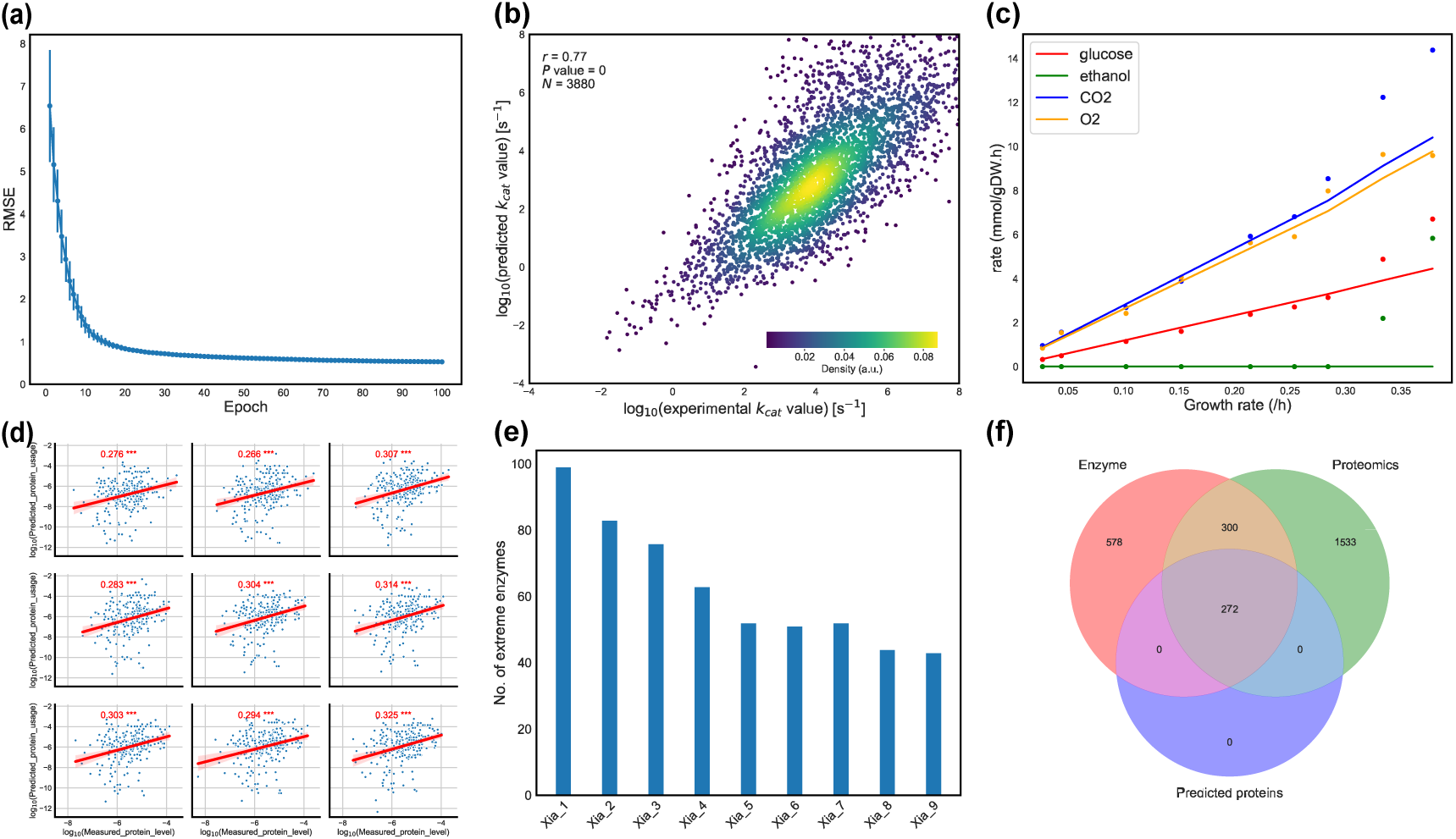
Bayesian training performance for ecGEM in predicting phenotypic traits and quantitative proteomic data. a. The RMSE during the Bayesian training process for phenotype measurement and prediction. Data represented the mean ± standard deviation (SD) of multiple replicates. b. Correlation between deep learning-predicted *kcat* values and *kcat* values optimized via the Bayesian approach. *P* value for the Pearson’s correlation was calculated using Student’s t-test. c. Predicted exchange rates for the Crabtree effect based on the optimized *kcat* values (lines) compared with experimental data (dots). d. Performance of ecGEM with optimized *kcat* values in predicting quantitative proteome. *: *P* value <0.05, **: *P* value <0.01, ***: *P* value <0.005. e. The number of extreme enzymes in the prediction of protein abundances. The extreme enzymes are defined as those with the ratio of predicted protein abundance to measured protein abundance more than 100 or less than 0.01. f. Venn diagram representing the overlap between measured proteins, predicted proteins, and enzymes included in the model.

**Supplemental Figure 2.**
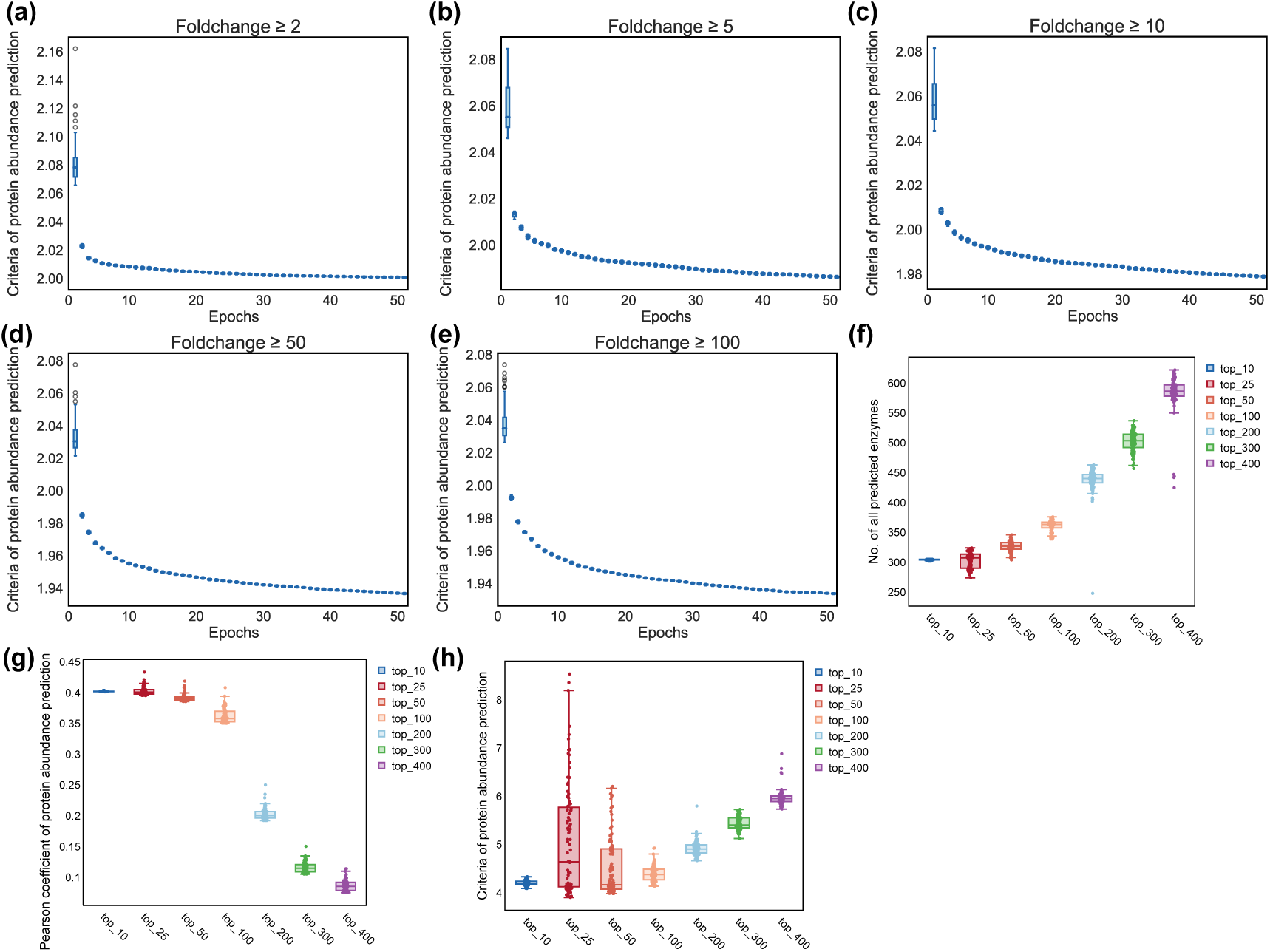
Comparison of optimization performance for extreme enzymes under different definitions. a-e. Performance of *kcat* optimization using the GAN method under different definitions of extreme enzymes, defined by fold change thresholds: more than 2 or less than 0.5 (a), more than 5 or less than 0.2 (b), more than 10 or less than 0.1 (c), more than 50 or less than 0.02 (d), and more than 100 or less than 0.01 (e). The training performance was represented by the absolute value of log2(fold change). f. Total number of predictable enzymes after optimization for each training process of the five definitions of extreme enzymes. g. The number of extreme enzymes identified after GAN-based optimization under each definition. h. Pearson correlation coefficient of protein abundance prediction after optimization, under each definition of extreme enzymes.

**Supplemental Figure 3.**
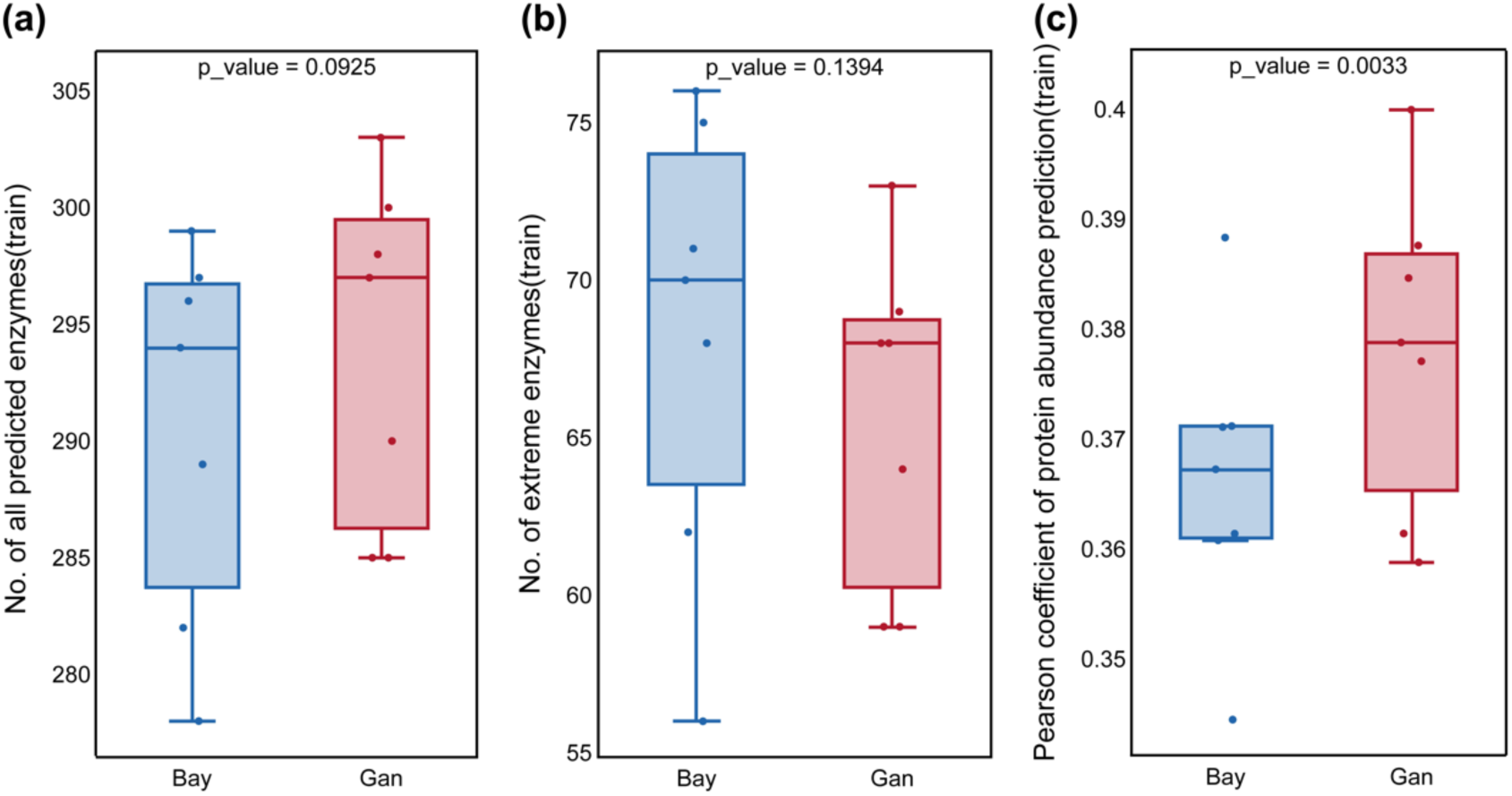
Performance of the GAN approach. a. Comparison of the total number of predictable enzymes using the GAN method versus the Bayesian approach. b. Comparison of the number of extreme enzymes using the GAN method and the Bayesian approach. c. Comparison of Pearson correlation coefficients of protein abundance predictions between the GAN method and the Bayesian approach.

**Supplemental Figure 4.**
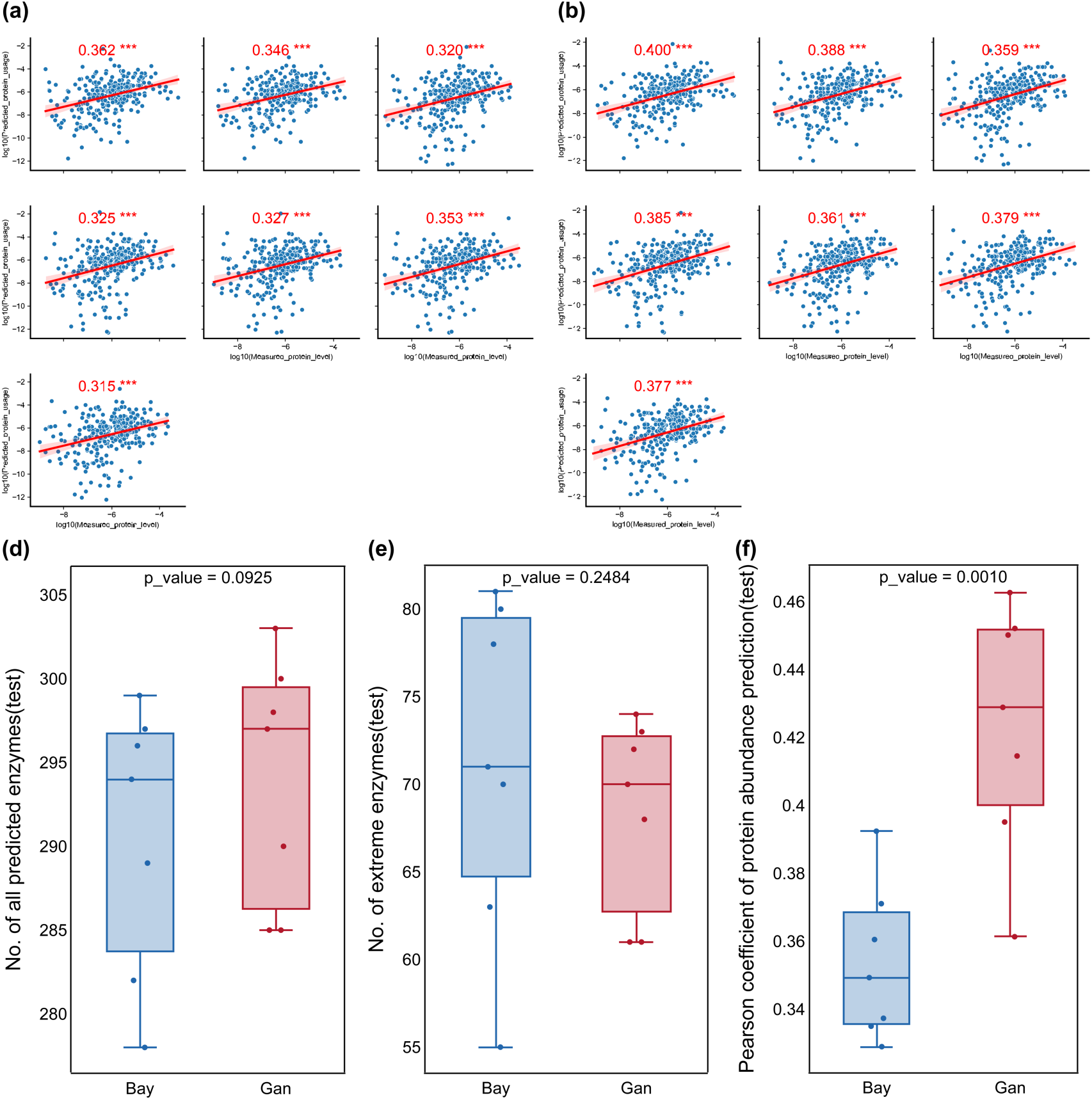
Performance of the GAN approach for protein abundance prediction under different nitrogen sources. a. Pearson correlation coefficients and corresponding *P* values for protein abundance predictions based on Bayesian training. *: *P* value <0.05, **: *P* value <0.01, ***: *P* value <0.005. b. Pearson correlation coefficients and corresponding *P* values for protein abundance predictions based on GAN training. *: *P* value <0.05, **: *P* value <0.01, ***: *P* value <0.005. c. Comparison of the total number of predictable enzymes under different nitrogen sources using the GAN method versus the Bayesian approach. d. Comparison of the number of extreme enzymes under different nitrogen sources using the GAN method and the Bayesian approach. e. Comparison of Pearson correlation coefficients of protein abundance predictions under different nitrogen sources between the GAN method and the Bayesian approach.

**Supplemental Figure 5.**
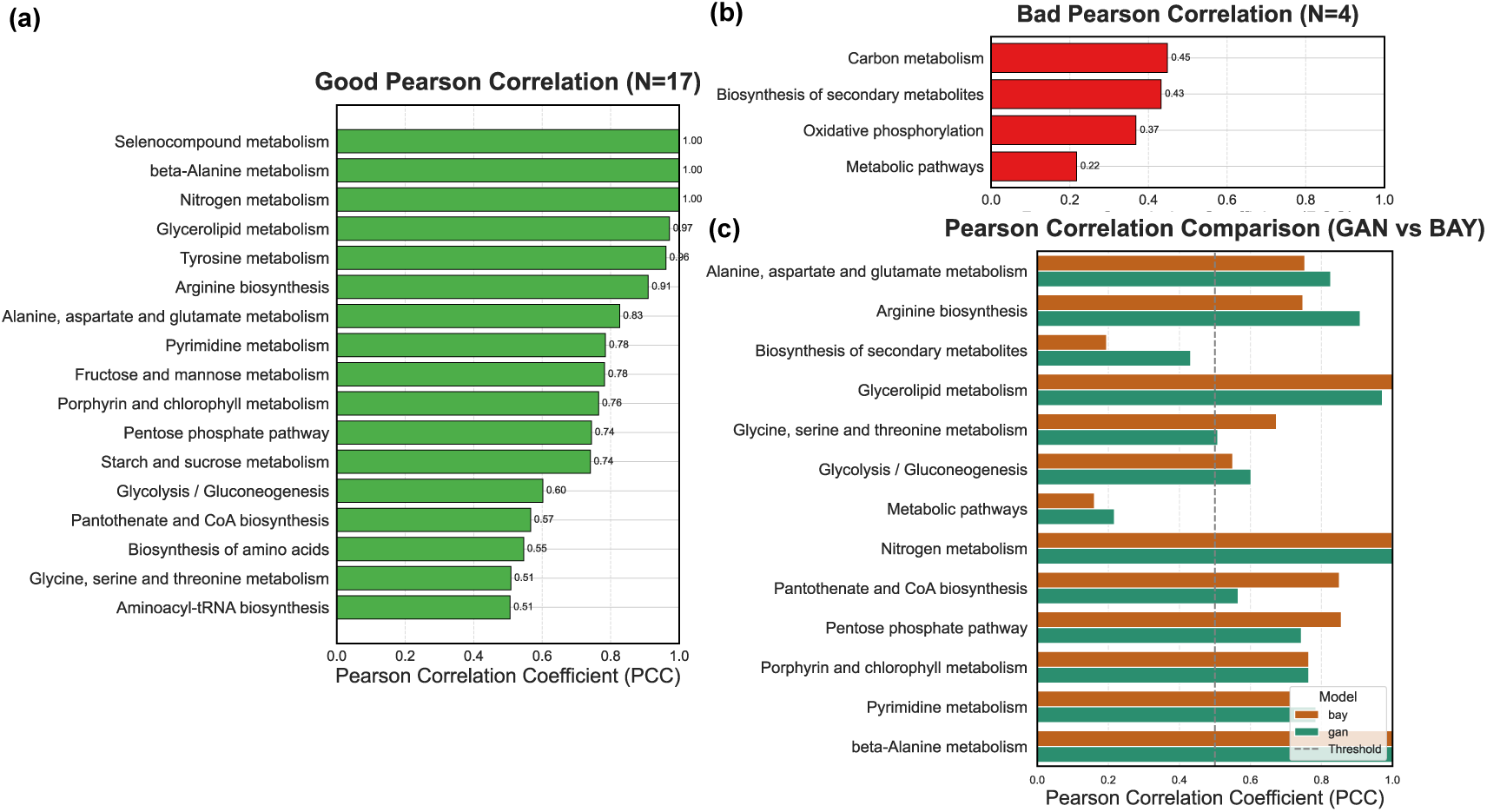
KEGG enrichment and method comparison of protein abundance prediction accuracy. a. KEGG pathways enriched among proteins with high prediction accuracy (Pearson correlation coefficients > 0.5) in the GAN-optimized ecGEM (GAN-ecGEM), along with their corresponding Pearson correlation coefficients (PCC). b. KEGG pathways associated with poorly predicted proteins (PCC < 0.5) in the GAN- ecGEM model. c. Comparison of PCC between GAN- and Bayesian (BAY)-optimized ecGEMs across shared proteins in the KEGG pathways.

**Supplemental Figure 6.**
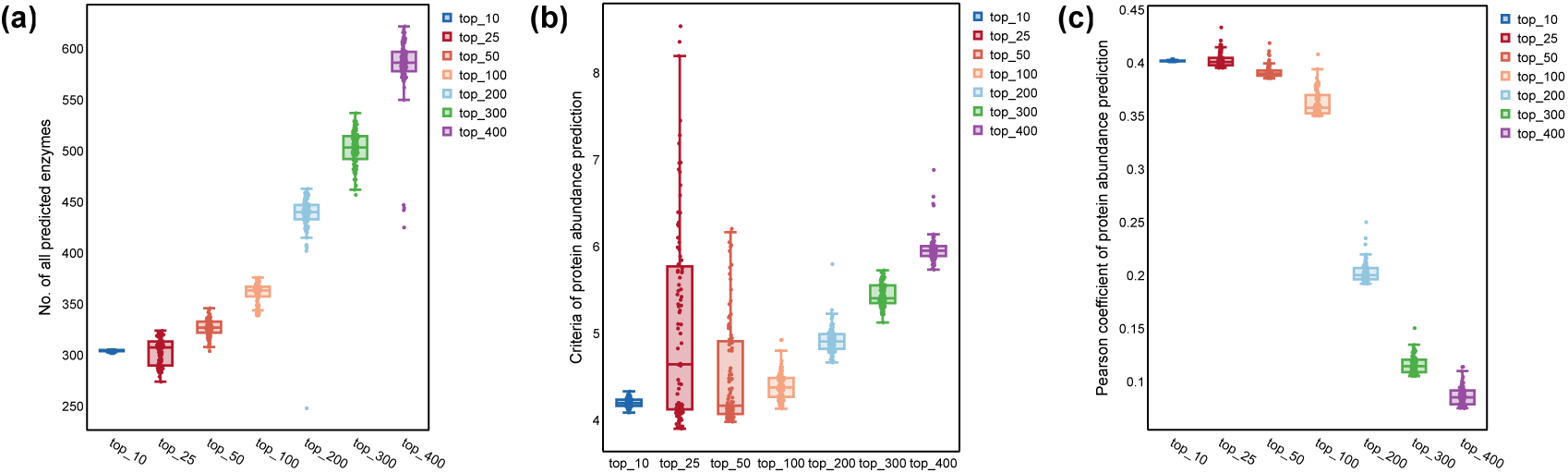
Comparison of the GAN approach in predicting protein synthetic rates at different scales a. Comparison of the total number of predictable enzymes across different scales of protein synthetic rate prediction using the GAN optimization approach. b. Comparison of the number of extreme enzymes under different scales of protein synthetic rate prediction with the GAN method. c. Comparison of Pearson correlation coefficients for protein abundance predictions across different scales.

**Supplemental Figure 7.**
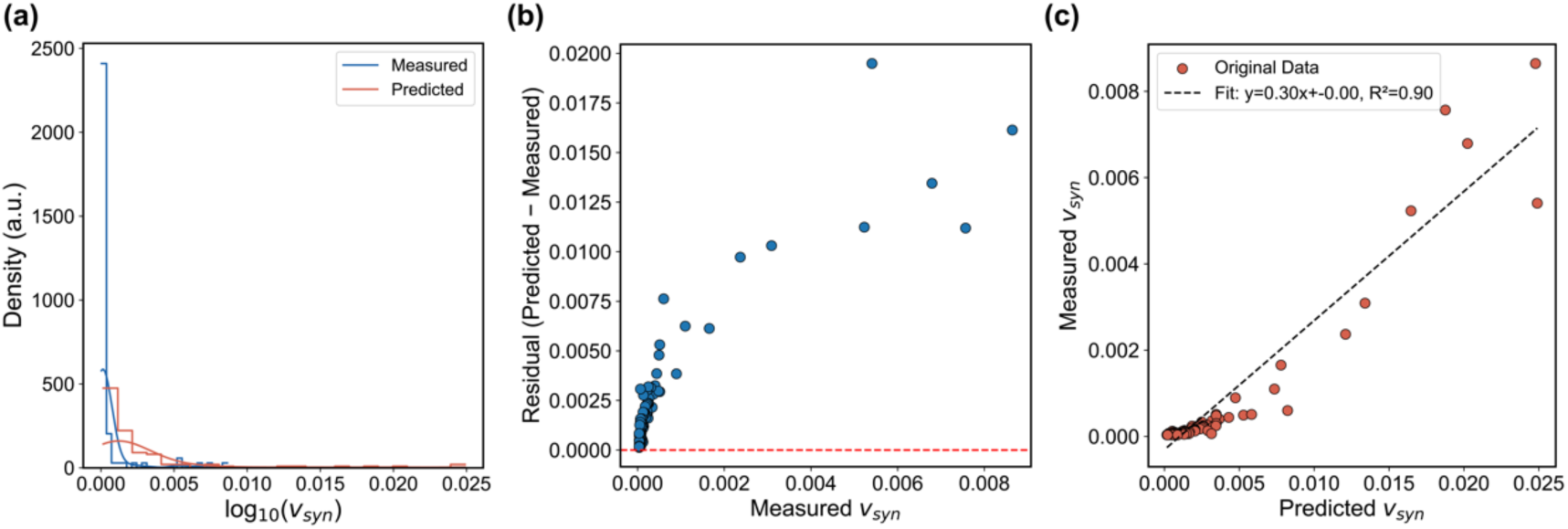
Evaluation of model performance for protein synthesis rate (*vsyn*) prediction. a. Density distribution of log10(*vsyn*) for predicted and measured values, showing overall consistency in trends. b. Residual plot (Predicted − Measured) against measured *vsyn* values. c. Scatter plot of predicted vs. measured *vsyn* values. The dashed line represents the linear regression fit.

